## Supplementary material for "Mouthparts of the bumblebee (*Bombus terrestris)* exhibit poor acuity for the detection of pesticides in nectar": suppplemental file

Supplemental Information

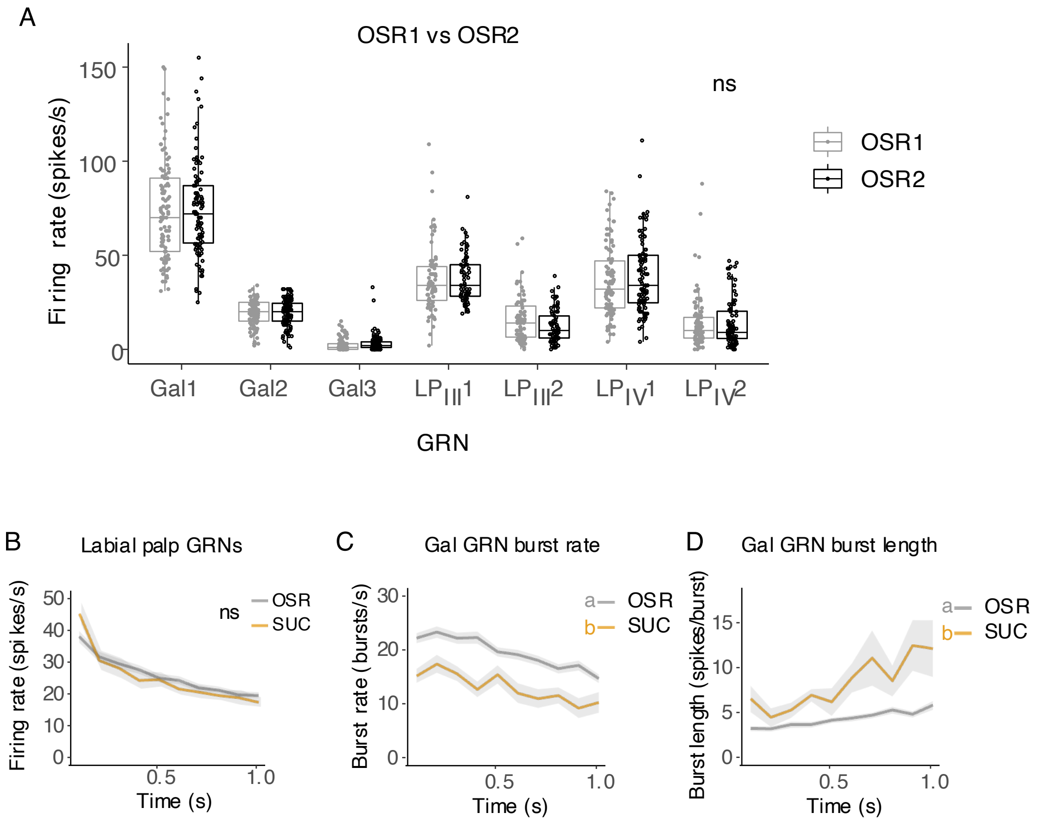


**Supplemental Information Figure 1: Electrophysiological responses to OSR**

A: Electrophysiological responses of GRNs were tested using a control solution of 10% OSR at the beginning and end of each session to ensure that the pesticides were not damaging the GRNs and that the responses remained stable. There were no differences in GRN responses to the first and last stimulations with OSR on any GRN (LME, stimulus: $F_{1,1267}$ = 0.1145, $p$ = 0.735, GRN: $F_{6,1267}$ = 569.98, $p$ < 0.0001).
B: Labial palp GRN firing rate histogram over 1 s stimulation with OSR or SUC with mean (line) and standard error of the mean (SEM) shading. There was no significant difference in labial palp GRN responses (n = 12 bees, LME, $F_{1,462}$ = 2.08, $p$ = 0.150).
C: Bursting rate histogram in 0.1 s bins following stimulation with OSR or SUC, with SEM in gray shading (n = 12 bees, LME, $F_{1,469}$ = 150, $p$ < 0.0001).
D: Average burst length (number of Gal1 spikes per burst) per 0.1 s bin following stimulation with OSR or SUC, with SEM in gray shading (n = 12 bees, LME, $F_{1,487}$ = 94.4, $p$ < 0.0001).

Parkinson et al, Bumblebee mouthparts exhibit poor acuity for the detection of pesticides in nectar

Supplemental Information

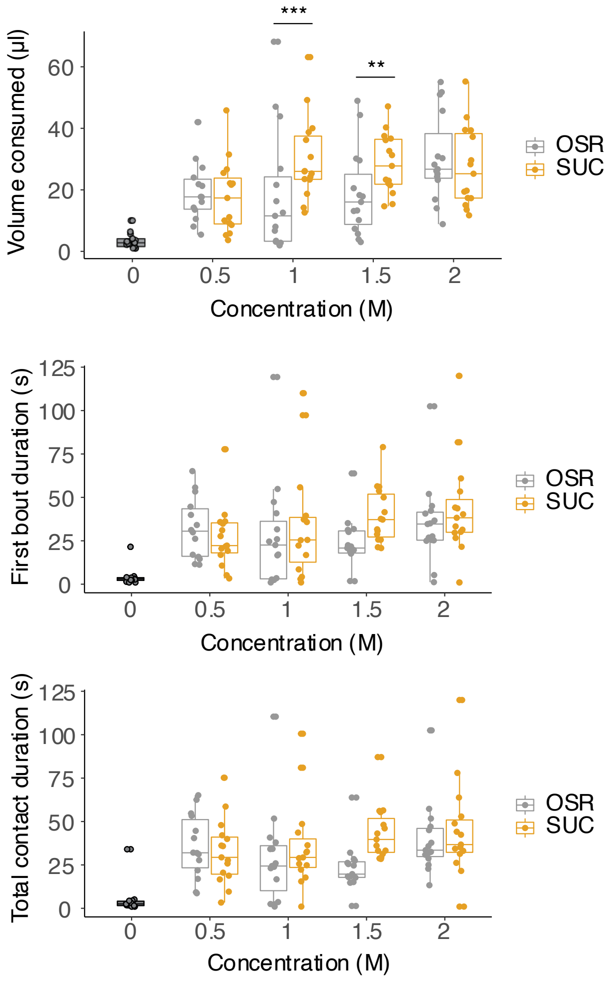


**Supplemental Information Figure 2: OSR and sucrose concentration gradients**
Feeding behaviour over 2 minutes was recorded using a range of sucrose and OSR concentrations (0, 0.5, 1.0, 1.5, and 2.0 M). Volume consumed (A, stimulus*concentration: $F_{4,133} = 4.51$, $p = 0.0019$), first bout duration (B, stimulus: $F_{1,136} = 2.05$, $p = 0.16$; concentration: $F_{4,138} = 27.5$, $p < 0.0001$), and cumulative contact duration (C, stimulus: $F_{1,137} = 2.20$, $p = 0.14$; concentration: $F_{4,138} = 40.4$, $p < 0.0001$) did not differ between stimuli, n=15 bees per stimulus.

Supplemental Information

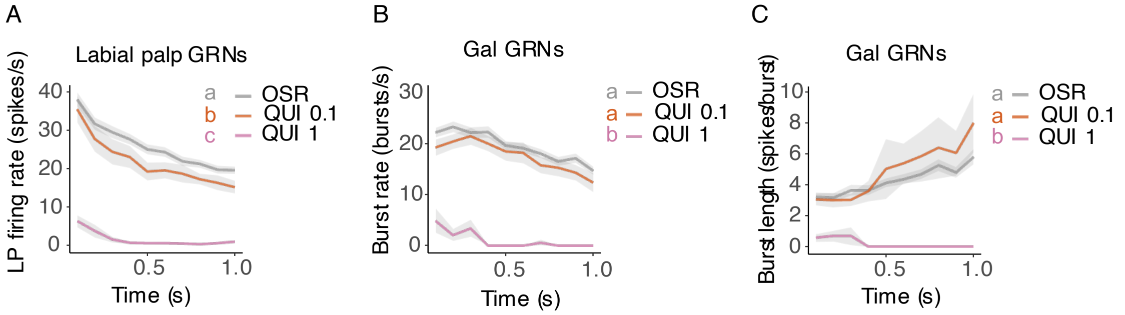


**Supplemental Information Figure 3: Temporal responses of GRNs to quinine**
A: Firing rate histogram showing the average firing rate of labial palp GRNs in 0.1 s bins following stimulation with 10% OSR, QUI 0.1 or QUI 1, with standard error of the mean (SEM) in gray shading (n = 12 bees, LME, stimulus $F_{2,554}$ = 576 $p$ < 0.0001). Results from estimated marginal means *post hoc* tests denoted by letters.
B: Bursting rate histogram in 0.1 s bins following stimulation with 10% OSR, QUI 0.1 or QUI 1, with SEM in gray shading (n = 12 bees, $F_{2,563}$ = 500, $p$ < 0.0001). Results from estimated marginal means *post hoc* tests denoted by letters.
C: Average burst length (number of Gal1 spikes per burst) per 0.1 s bin following stimulation with 10% OSR, QUI 0.1 or QUI 1, with SEM in gray shading (n = 12 bees, LME $F_{2,574}$ = 97.9, $p$ < 0.0001). Results from estimated marginal means *post hoc* tests denoted by letters.

Supplemental Information

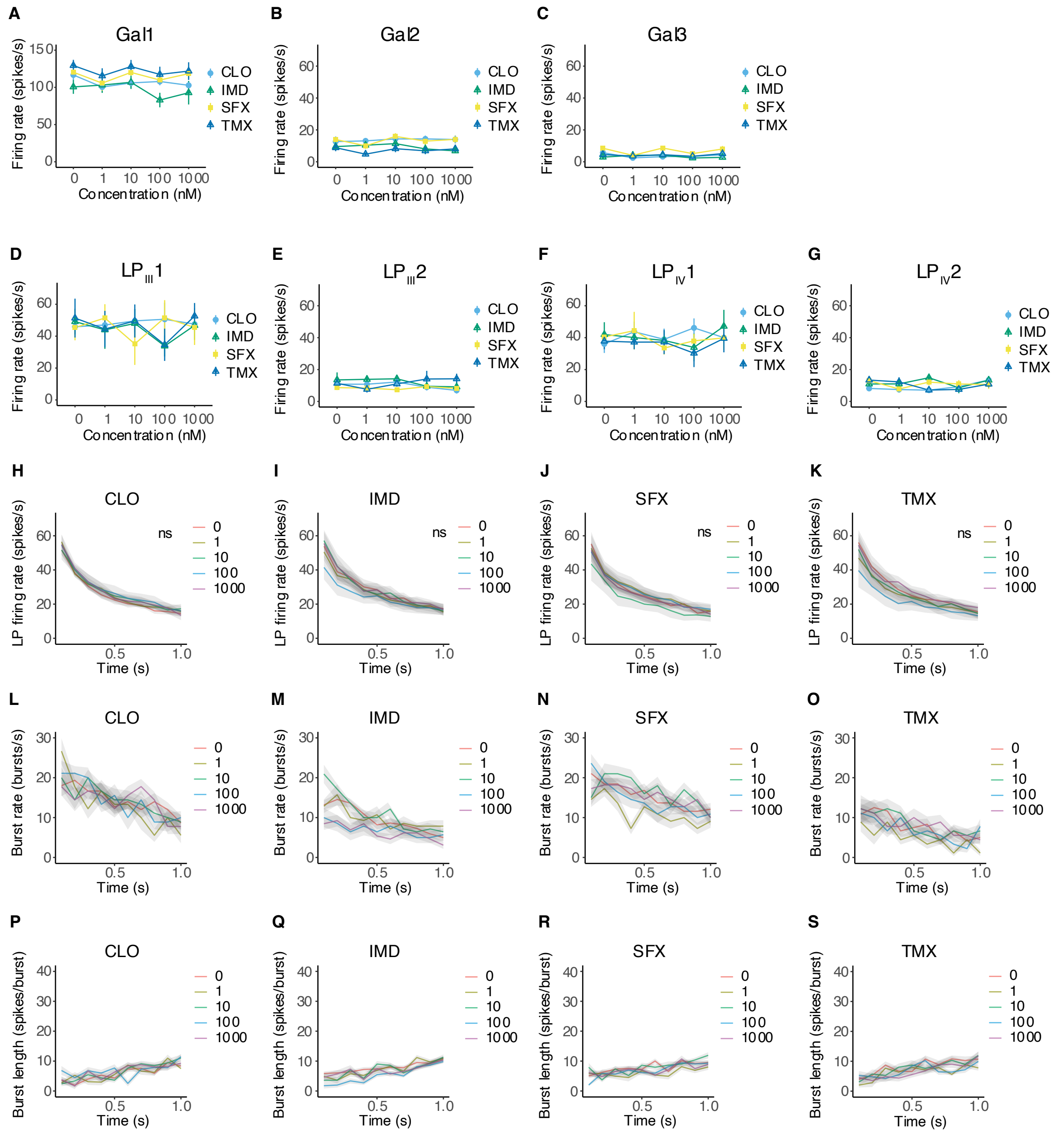


**Supplemental Information Figure 4: Galeal and labial palp GRN cannot detect low concentrations of pesticides in nectar.**

A-G: Average firing rate over 1s of galeal and labial palp GRNs across a concentration gradient of pesticides mixed with a 10% dilution of OSR. Galeal GRN1 plots show mean + standard error. There was no significant effect of concentration on GRN firing across locations ($F_{1,2727}$ = 0.51, $p$ = 0.48, n=18-28 sensilla per stimulus) while there was a significant interaction between stimulus type and GRN ($F_{18,2716}$ = 8.22, $p$ < 0.0001). *Post hoc* estimated marginal means revealed significant differences of galeal GRN2 and GRN3 firing rates between stimuli.

Parkinson et al, Bumblebee mouthparts exhibit poor acuity for the detection of pesticides in nectar

Supplemental Information

H-K: Firing rate histograms of labial palp GRNs in 100 ms bins over 1 s of stimuluation with CLO (H, concentration: $F_{1,832}$ = 1.95, $p$ = 0.16; time: $F_{1,832}$ = 0.11, $p$ = 0.74, n=18), IMD (I, concentration: $F_{1,823}$ = 3.55, $p$ = 0.064; time: $F_{1,823}$ = 696.67, $p$ < 0.0001, n=28), SFX (J, concentration: $F_{1,863}$ = 0.51, $p$ = 0.48; time: $F_{1,863}$ = 645.44, $p$ < 0.0001, n=22) TMX (K, concentration: $F_{1,833}$ = 0.32, $p$ = 0.57; time: $F_{1,833}$ = 635.30, n=18).
L-O: Burst rate (GRN2 rate) histograms over 1s of stimulation were not affected by CLO (L, concentration: $F_{1,832}$ = 1.947, $p$ = 0.163; time: $F_{1,832}$ = 493.36, $p$ < 0.0001, n=18), IMD (M, concentration: $F_{1,823}$ = 3.435, $p$ = 0.0642; time: $F_{1,823}$ = 696.67, $p$ < 0.0001, n=28), SFX (N, concentration: $F_{1,863}$ = 0.507, $p$ = 0.477; time: $F_{1,823}$ = 645.44, $p$ < 0.0001, n=22), and TMX (O, concentration: $F_{1,833}$ = 0.318, $p$ = 0.573; time: $F_{1,833}$ = 635.03, $p$ < 0.0001, n=18).
P-S: The number of GRN1 spikes per burst was affected by imidacloprid but not other pesticides (P, CLO, concentration: $F_{1,1011}$ = 3.05, $p$ = 0.0811; time: $F_{1,1011}$ = 6.99, $p$ = 0.0083, n=18, Q, IMD, concentration: $F_{1,1529}$ = 12.82, $p$ = 0.00035; time: $F_{1,1529}$ = 33.88, $p$ < 0.0001, n=28, R, SFX, concentration: $F_{1,1231}$ = 1.83, $p$ = 0.18; time: $F_{1,1231}$ = 0.47, $p$ = 0.0012, n=22, S, TMX, concentration: $F_{1,1231}$ = 1.83, $p$ = 0.18; time: $F_{1,1231}$ = 0.47, $p$ = 0.0012, n=18).

Parkinson et al, Bumblebee mouthparts exhibit poor acuity for the detection of pesticides in nectar

Supplemental Information

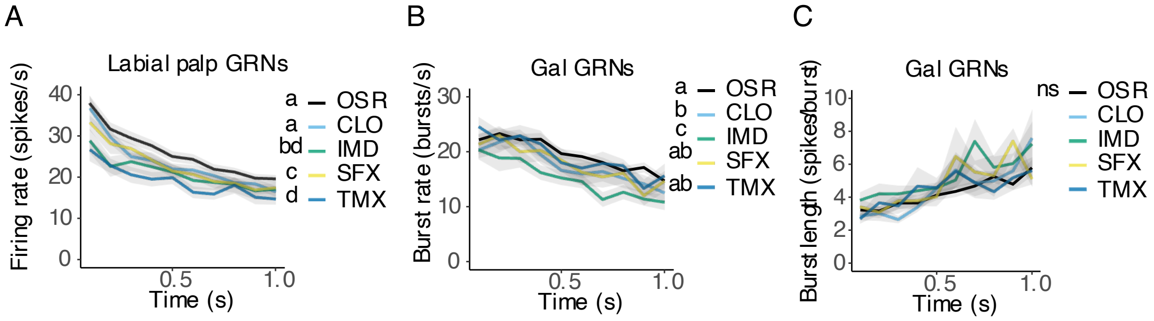


**Supplemental Information Figure 5: Temporal responses of GRNs to high concentrations of pesticides**

A: Firing rate histogram showing the average firing rate of labial palp GRNs in 0.1 s bins following stimulation with OSR, CLO, IMD, SFX, or TMX, with standard error of the mean (SEM) in gray shading (n = 12 bees, LME, stimulus: $F_{4,817} = 44.2$, $p < 0.0001$; time: $F_{1,808} = 884$, $p < 0.0001$). Results from estimated marginal means *post hoc* tests denoted by letters.

B: Bursting rate histogram in 0.1 s bins following stimulation with OSR, CLO, IMD, SFX, or TMX, with SEM in gray shading (n = 12 bees, LME, stimulus: $F_{4,825} = 13.3$, $p < 0.0001$; bin: $F_{1,808} = 401$, $p < 0.0001$). Results from estimated marginal means *post hoc* tests denoted by letters.

C: Average burst length (number of Gal1 spikes per burst) per 0.1 s bin following stimulation with OSR, CLO, IMD, SFX, or TMX, with SEM in gray shading (n = 12 bees, LME, stimulus: $F_{4,841} = 1.69$, $p = 0.150$; time: $F_{4,841} = 1.69$, $p = 0.150$). Results from estimated marginal means *post hoc* tests denoted by letters.
